# Inducible nitric oxide synthase (NOS2) knockout mice as a model of trichotillomania

**DOI:** 10.1101/086173

**Authors:** PC Casarotto, C Biojone, K Montezuma, FQ Cunha, SRL Joca, E Castren, FS Guimaraes

## Abstract

Trichotillomania (TTM) is an impulse control disorder characterized by repetitive hair pulling/trimming. Barbering behavior (BB) has been observed in laboratory animals and proposed as TTM model. The neurobiological basis of TTM is not clear, but it seems to involve striatal hyperactivity parallel to hypoactivation of prefrontal cortex. In this study we observed that knockout mice to the inducible isoform of nitric oxide synthase (NOS2) exhibit exacerbated BB following the 4th week of age, as well as increased repetitive movements compared to wild-type mice (WT). These behaviors are associated to decreased levels of NMDA receptor subunit (NR1) in prefrontal cortex, while an increase was observed in striatum of NOS2KO compared to WT. Striatal neurons from NOS2KO also exhibited increased number of branches compared to WT. The repeated treatment with clomipramine, a clinically approved drug to treat TTM in humans, or memantine, an antagonist of NMDA receptors, as well as partial rescue of NOS2 expression in haploinsufficient animals, attenuated the expression of BB. The silencing of NOS2 expression reduced the MAP2 (microtubule-associated protein 2) levels in activity-induced differentiated PC12 cells. Our data led us to propose that NOS2 regulates the neuronal maturation of the inhibitory afferent pathways to striatum during neurodevelopment, and such inadequate inhibition of striatal motor programs might be associated to the observed phenotype.

## Introduction

The cortico-striato-thalamo-cortical circuitry (CSTC) is a series of reverberatory loops that control motor programs. The glutamatergic neurotransmission plays a crucial role in both afferent and efferent pathways in prefrontal cortex and striatum, two core structures in the regulation of CSTC (Langen et al., 2011a, 2011b). In this scenario, nitric oxide (NO), plays a major part, not only regulating the release of glutamate but also in the migration and maturation of neuronal cells in cortex. This later feature of NO is apparently mediated by the inducible isoform of nitric oxide synthase (NOS2) (Arnhold et al., 2002).

Malfunctions of specific parts of CSTC is speculated to be involved in the neurobiology of several motor and impulse control disorders, such as obsessive-compulsive disorder, Tourette's syndrome, and, of interest to the present study, trichotillomania (Langen et al., 2011a, 2011b).

Trichotillomania (TTM) is an impulse control disorder characterized by a distressing urge for hair pulling or trimming, affecting 1-3.5% of the population (American Psychitaric Association, 2016; Christenson et al., 1991). Nowadays the pharmacotherapy for TTM is based on treatment with the serotonin reuptake inhibitor clomipramine (McGuire et al., 2014; Swedo et al., 1993, 1989). Recent studies also indicate that the glutamatergic neurotransmission modulator n-acetylcysteine, is effective in alleviating the symptoms of TTM (Grant et al., 2009). Other drugs, such as atypical antipsychotic olanzapine, have shown success in placebo-controlled trial (Van Ameringen et al., 2010).

Neuroimaging studies indicated increased gray matter volume in striatum, amygdalo-hippocampal formation, and multiple cortical regions (including the cingulate, supplemental motor cortex, and frontal cortex) of patients with TTM (Chamberlain et al.,2009). This increase in gray matter was associated with a relative decreased activity in cortical structures, and hyperactivity in the striatal region (Fineberg et al., 2010). Despite these findings, the neurobiology of TTM is not completely understood.

Animal models for the investigation of neurobiological aspects of TTM as well as the search for new therapies rely on the observation of barbering behaviors (BB). In laboratory and captive animals, BB is characterized by pulling or trimming of fur or whiskers (Garner et al., 2004). Some transgenic strains, e.g. SAPAP3 and SLITRK5 knockouts, display compulsive/repetitive behavior (Shmelkov et al., 2010; Welch et al., 2007), however no specific BB has been described to them. Interestingly, SAPAP3 is a structural protein crucial for a properly docking of glutamatergic receptors (Welch et al., 2007).

In the present study we describe the BB observed in NOS2KO mice, associated to alterations in the prefrontal cortex and striatum. We also assessed the effects of chronic treatment with clomipramine, or memantine (a non-competitive antagonist of NMDA receptors), and the partial rescue of NOS2 expression on BB.

## Experimental procedure

### Animals

Male and female NOS2 knockout (NOS2KO; Jackson Lab #2596) and C57BL6/j (WT) as controls were used, with food and water available *ad libitum* except during experimental procedures. All efforts were made to minimize animal suffering. All protocols were approved and are in accordance with international guidelines for animal experimentation.

### Drugs

Clomipramine hydrochloride (clomipramine; Sigma-Aldrich, #C7291) and memantine hydrochloride (memantine; Sigma-Aldrich, #M9292) were used. All drugs were dissolved in the drinking water. The solutions were changed every 2 days, and the amount of liquid consumed was determined by changes in the bottle weight. The total volume was divided by the number of animals in each cage and an average of 5ml/day/animal was stably observed throughout the experiments.

### Actimeter test (ACTM)

The actimeter test was performed to evaluate general locomotor activity as well as repetitive behaviors. This apparatus consisted of two frames of 32 infrared beams in a square arena (Panlab-Harvard Apparatus, Barcelona, Spain). The animals were allowed to explore the arena for 15min.

### Barbering behavior (BB)

The animals were scored regard the state of the whiskers and fur, hair trimming or plucking. The score (table 1) was based on Garner and colleagues (Garner et al., 2004) body map for BB. The sum score for the absence of whiskers/fur in snout, eye surroundings, forehead, chest, neck and back was determined every 5 days by an observer blind to the drug treatment.

**Table 1.** Score for barbering behavior. Based on Garner et al 2004.

| Area affected | score |
| --- | --- |
| No hair loss | 0 |
| Loss of whiskers | +1 |
| Hair loss in snout | +1 |
| Hair loss in eye area | +1 |
| Hair loss in forehead | +1 |
| Hair loss in chest/neck | +1 |
| Hair loss in back | +1 |

### Sample collection and western blotting

Animals were deeply anesthetized with 4% chloral hydrate injected intraperitoneally, the brains were removed and dissected on ice. Samples from sriatum (STR) and prefrontal cortex (PFC) were mechanically homogenized in lysis buffer [137mM NaCl; 20mM Tris-HCl; 10% glycerol) containing protease/phosphatase inhibitor cocktail (Sigma-Aldrich, #P2714; #P0044) and sodium orthovanadate (0.05%, Sigma-Aldrich, #S6508)]. The homogenate was centrifuged (10000 x g) at 4^o^C for 15min and the supernatant collected and stored at -80°C. Thirty micrograms of total protein content in each sample was separated in acrylamide gel electrophoresis and transferred to a PVDF membrane. Following blockade with 5%BSA in TBST (20mM Tris-HCl; 150mM NaCl; 0.1% Tween20) the membranes were exposed to primary antibody against subunit NR1 of NMDA receptor (Cell Signaling, USA, #5704) or GAPDH (Santa Cruz, USA, #sc25778) overnight at 4°C. After washing with TBST the membranes were incubated with anti-rabbit IgG HRP-conjugated antibody 2h at RT. Following washing the membranes were incubated with 4-chloronaphtol (4CN, Perkin Elmer, #NEL300001EA) for color development. The dried membranes were scanned and the intensity of bands determined using ImageJ software (NIH, v.1.49).

### Golgi method for silver impregnation

Brains from perfused mice were treated according to manufacturer’s instructions (FD Neurotechnologies, #PK401, USA). Briefly, the brains were incubated in solution A+B for 14 days, then transferred to solution C for 72 hours. Following the incubation in solution C, the brains were cut in a cryostat (sections of 100 micrometers – um). The coronal sections were transferred to gelating-coated microscope glass slides, allowed to dry and photographed at 100x magnification in a light microscope. The number of neuronal branches was were determined along the neurites by Sholl analysis using ImageJ software (NIH, v.1.49).

### Cell culture, transfection, differentiation and determination of MAP2 levels

PC12 cells, clone 615, were maintained in DMEM medium, supplemented with 10% fetal bovine serum (FBS) and 5% horse serum (HS), containing penicillin/streptomycin and glutamine. For the experiments, the cells were transferred to 12-well plates, and 24 after, the cultivation medium was changed to DMEM 0.5%FBS at a density of 5x10^5^cells/well. Twenty-four hours later, the cells were transfected with a mixture of lipofectamine 2.5% in OptiMEM medium and 20ug of plasmid containing 4 different interference sequences (shRNA) for NOS2 (5ug of each sequence, #TR709254). Twenty-four hours after transfection, the cultivation medium was changed and the cells were incubated with KCl (25mM) for 5days. The levels of microtubule-associated protein 2 (MAP2) were determined by direct ELISA. Briefly, the samples (120ug of total proteins) were incubated in transparent 96-well U shaped plates overnight at 4°C. Following blockade with 1% BSA in PBST, antibody against MAP2 (Millipore, #AB5622, USA) was incubated overnight at 4°C. The development of color following incubation with anti-rabbit IgG-HRP and BM Blue POD substrate (Roche, #11484281001, USA) was stopped by 1M of HCl and the color intensity was read at 450nm. The sample read outs, discounted the blank values, were expressed as percentage from control-group (scrambled).

## Experimental procedures

### Experiment 1: time course of drug treatment effect in BB

NOS2KO male and female mice (8-9 weeks of age at the beginning of the experiment) were scored for BB and received clomipramine (10mg/kg) for 20 days in the drinking water. The average amount of solution consumed by the animals was monitored every 2 days to correct the drug intake. A separated cohort of female NOS2KO mice received memantine (1mg/kg) for 20 days in the drinking water. The BB was scored at the start of the treatment and monitored every 5days until the end of the experiment.

### Experiment 2: repetitive behavior in NOS2KO and WT mice

NOS2KO and WT were submitted to the actimeter for 15min. The total number of interruptions of the infrared beams was considered as total activity (Casarejos et al., 2013). The number of repetitive movements (including stereotypies and grooming) were also counted by the apparatus and normalized by the total activity, therefore expressed as percentage of repetitive behavior.

### Experiment 3: effect of clomipramine in NOS2KO repetitive behavior

NOS2KO received clomipramine (approximately 10mg/kg/day, in the drinking water) or vehicle during 14 days. On day 13, half of the vehicle treated group received clomipramine in the drinking water. Therefore the following groups were obtained: vehicle, acute clomipramine, and repeated clomipramine.

### Experiment 4: NR1 expression and evaluation of neuronal branches in prefrontal cortex and striatum of NOSKO and WT

NOS2KO and WT mice were euthanized, as described above, and samples from PFC and striatum were assay for the levels of NMDA receptor subunit 1A (NR1). A separated cohort of NOS2KO and WT animals was anesthetized, perfused and the brains collected for silver impregnation by Golgi’s method.

### Experiment 5: rescue of BB by partial expression of NOS2

NOS2.haploinsufficient animals were obtained y crossing of NOS2KO males and WT females. The level of BB in the offspring was observed from 2nd to 9th week of age. Offspring from NOS2KO parents born in the same periods were used as controls.

### Experiment 6: role for NOS2 on activity-induced expression of MAP2 in PC12 cells

PC12 cells, transfected with NOS2 shRNA or scramble (non-coding sequence) were differentiated in 25mM KCl for 5 days. At day 5 the cells were lysed and the levels of MAP2 determined by ELISA.

## Statistical analysis

The data from experiments 1, 4 (silver impregnation) and 5 was analyzed by two-way ANOVA (considering drug treatment or genotype, and time or distance from soma as factors). The data from experiments 2, 4 (western-blotting) and 6 was analyzed by Student’s t test. Finally, the data from exp 3 was analyzed by one-way ANOVA. The Fisher’s LSD *post hoc* test was used when appropriate. Unless otherwise stated all data is expressed as Mean/SEM. Values of p<0.05 were set as significant.

## Results

### Experiment 1: time course of drug treatment effect in BB

As shown in fig 1b–d, there is a significant effect of clomipramine treatment on the BB in both males [F(1,18)= 27.0, p<0.05] and females [F(1,8)= 29.04, p<0.05]. The *post hoc* analysis indicates a significant difference between clomipramine and vehicle treated groups at days 10th, 15th and 20th for both males and females (Fisher’s LSD, p<0.05). The treatment with memantine was also effective on BB [F(1,12)= 38.69, p<0.05], with significant difference from controls observed at days 10th, 15th and 20th.

**Figure 1.**
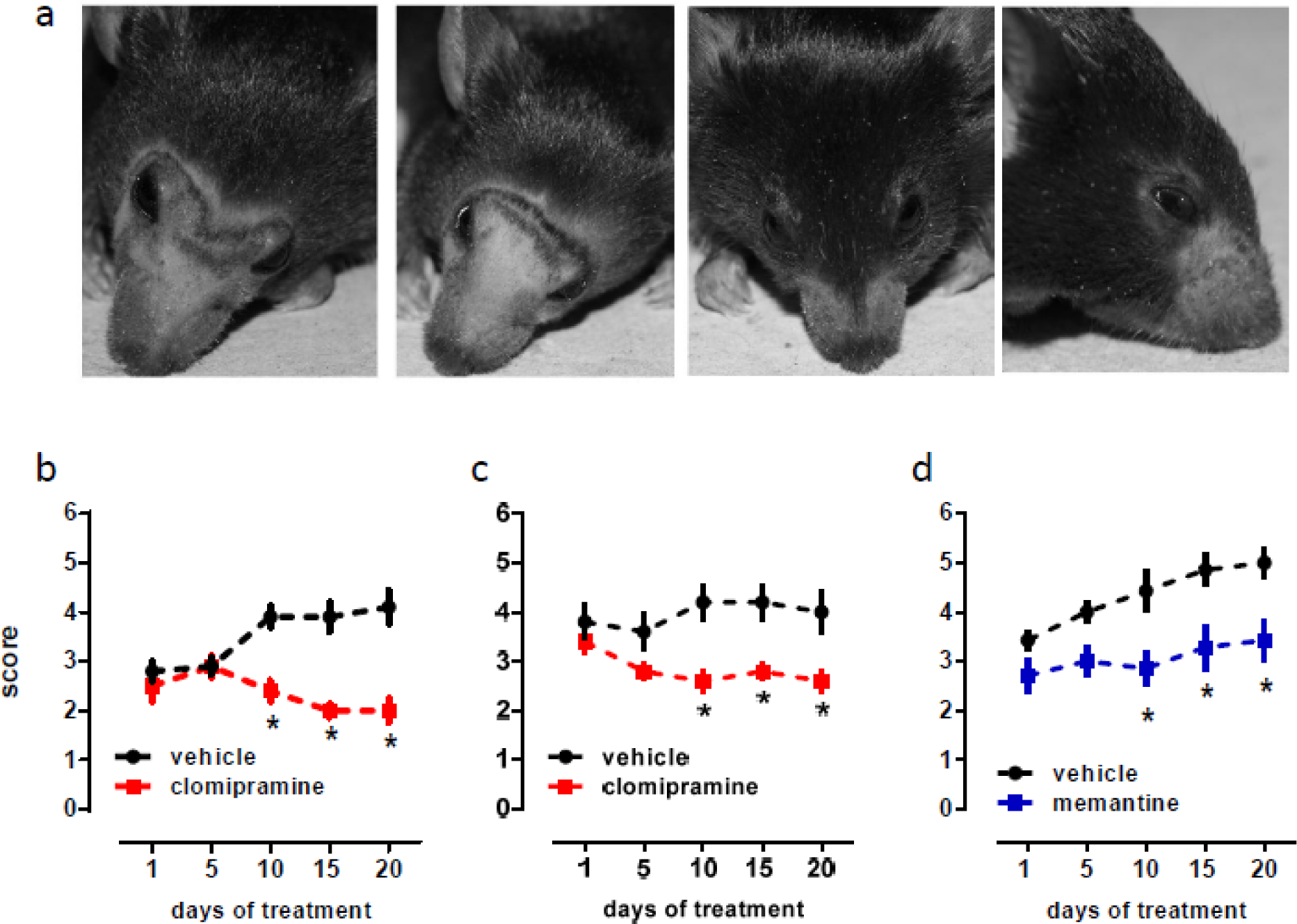
(a) phenotype of NOS2KO barbering behavior (BB), showing two vehicle- (left panels) and two clomipramine-treated (right pannels) female NOS2KO. Effect of clomipramine (red squares; 10mg/kg in the drinking water) on BB in males (b, n=10/group) and females (c; n=5/group) NOS2KO mice. (d) effect of memantine (blue squares; 1mg/kg in the drinking water) on BB in NOS2KO female mice (n=7/group). *p<0.05 from vehicle group at the same time point.

### Experiment 2: repetitive behaviors in NOS2KO and WT mice

The Student’s t test indicates a significant effect of genotype, with NOS2.KO exhibiting reduced total activity in males [t(10)= 2.28, p<0.05] and females [t(11)= 4.79, p<0.05], fig 2a. An increase in the percentage of repetitive movements was observed [males: t(10)= 4.34; females: t(11)= 2.69, p<0.05 for both], as found in fig 2b.

**Figure 2.**
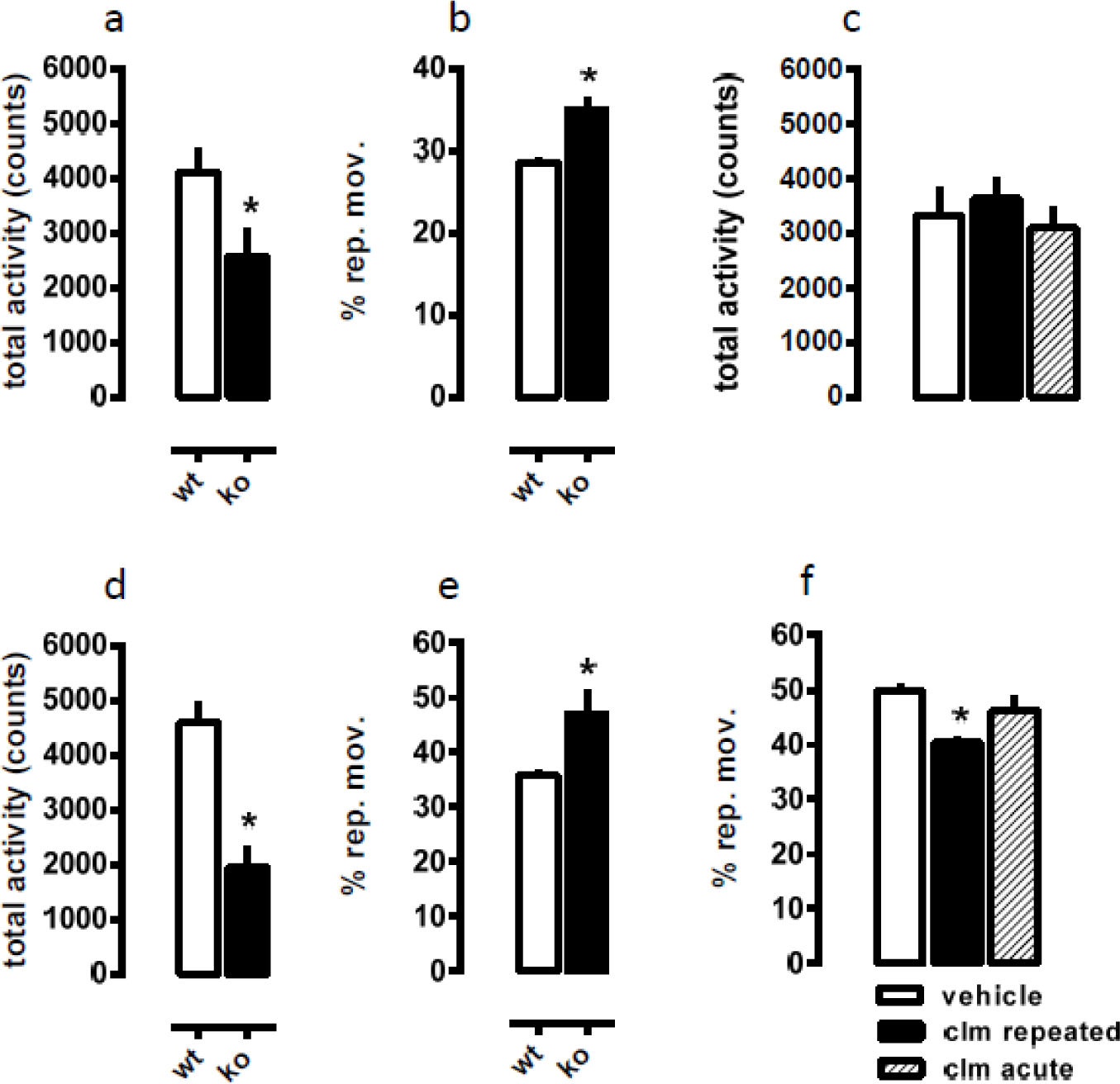
Reduced total activity and increased percentage of repetitive behaviors in NOS2KO males (a-b; n= 6-7/group) and females (d-e; n= 6-7/group) compared to WT. Effect of clomipramine (10mg/kg in the drinking water) on total activity (c) and repetitive behaviors (f) in female NOS2KO (n= 5-6/group). *p<0.05 from WT or vehicle-treated group.

### Experiment 3: effect of clomipramine in NOS2KO repetitive behavior

As observed in fig 2c, there was no significant effect of clomipramine treatment on total activity in NOS2KO [F(2,14)= 0.43, p>0.05]. However, repeated, but not acute, treatment with clomipramine decreased the percentage of repetitive movements compared to the control group [F(2,14)= 6.18, p<0.05], as found in fig 2f.

### Experiment 4: NR1 expression and evaluation of neuronal branches in prefrontal cortex and striatum of NOSKO and WT

The analysis by Student’s t test indicate a significant effect of genotype decreasing the levels of NR1 in PFC [t(10)= 3.08, p<0.05] and increasing NR1 in striatum [t(10)= 2.24, p<0.05], as seen in fig 3a–b, respectively.

**Figure 3.**
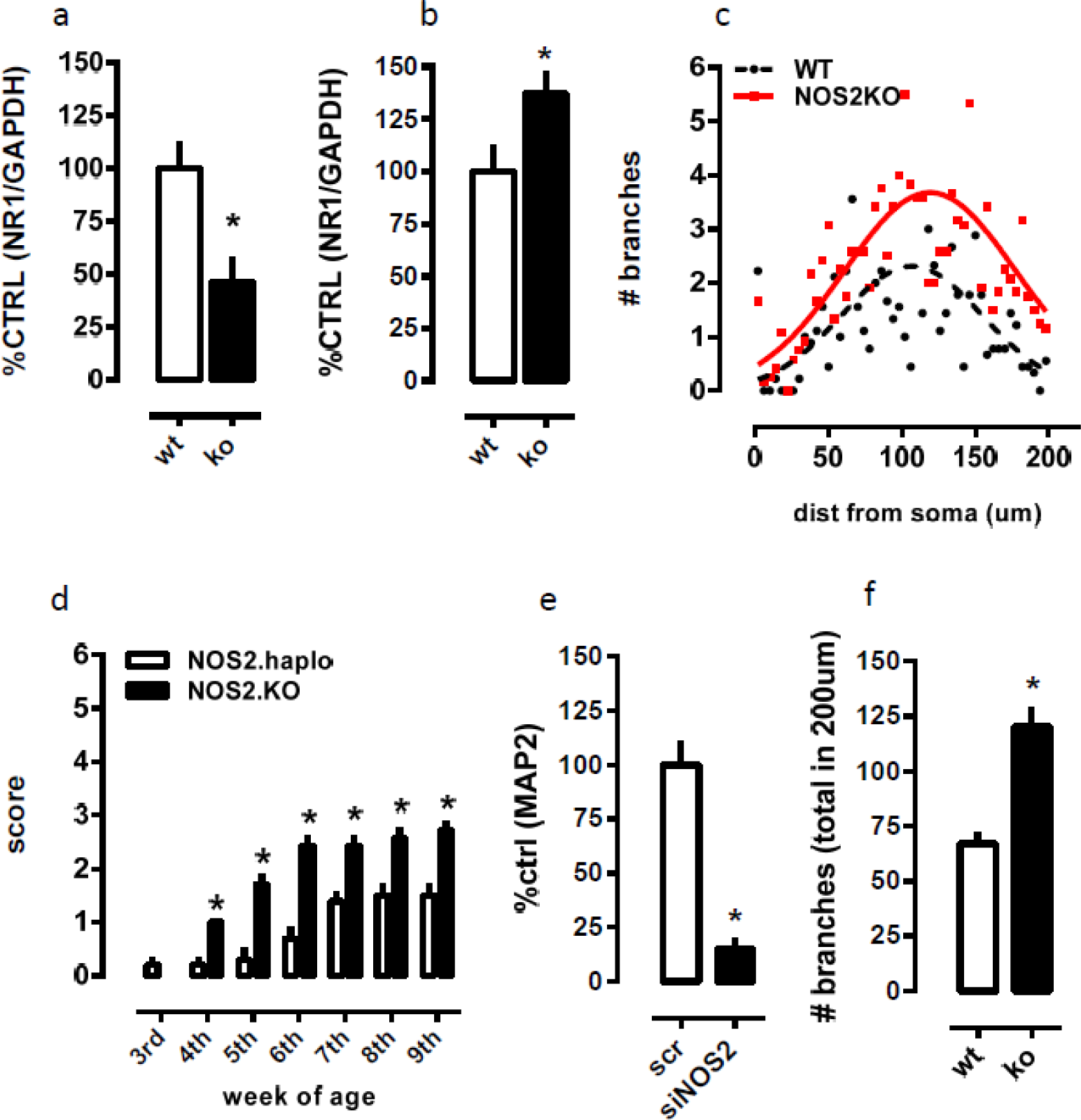
Changes in the levels of NR1 subunit in prefrontal cortex (a) and striatum (b) of female NOS2KO vs WT (n=6/group). (c and f) increase in the number of branches in the striatum of NOS2KO (red, continuous line) vs WT (black dashed line; n= 9-12 cells / group). (d) the partial rescue of NOS2 expression delays the onset and intensity of barbering behavior (n= 7-10/group). (e) the silencing of NOS2 compromises the expression of MAP2 expression in KCl-induced differentiation of PC12 cells. *p<0.05 from WT, NOS2 haploinsufficient or scrambled (scr) groups.

The analysis by two-way ANOVA indicates a significant effect of the genotype [F(1,19)= 20.41, p<0.05] and interaction [F(49,931)= 2.77, p<0.05] increasing the number of branches in coronal slices of striatum from NOS2KO compared to WT distributed alongside 200um from the soma, as found in fig 3c. The total number of branches was also found increased in NOS2.KO compared to WT [t(19)= 4.52, p<0.05], fig 3f.

### Experiment 5: rescue of BB by partial expression of NOS2

The analysis by two-way ANOVA indicates a significant effects of the genotype [F(1,105)= 99.87, p<0.05], the age [F(6,105)= 34.14, p<0.05] and interaction between these factors [F(6.105)= 5.23, p<0.05] in the barbering behavior exhibited by NOS2KO compared to NOS2 haploinsufficient animals from weaning age, fig 3d. The Fisher’s LSD test indicate significant differences starting from the 4th until 9th week of age (Fisher’s LSD, p<0.05). The design using NOS2KO sire and WT mother was chosen due to the allobarbering displayed by NOS2KO, so using WT females it was possible to avoid any effect of mother’s BB on pups, however no difference was observed at the weaning age (3rd week) between pups from WT and NOS2KO mothers.

### Experiment 6: role for NOS2 on activity-induced expression of MAP2 in PC12 cells

The Student’s t test indicates a significant effect of NOS2 silencing on the levels of MAP2 in PC12 cells [t(7)= 6.60, p<0.05], as found in fig 3e.

## Discussion

The present study suggests that the increased barbering behavior (BB) of NOS2KO mice as could be a putative model of trichotillomania (TTM). NOS2KO exhibited increased BB after weaning (significantly higher at 4th week of age), and this phenotype increases until adulthood. TTM is an impulse control disorder, characterized by hair pulling or trimming. To our knowledge, there is no established animal model to study this disorder, which reflects in the poor pharmacotherapy and interventions available (Johnson and El-Alfy, 2016). In animals BB is a rare phenotype, with an incidence of approximately 2.5% in laboratory mice (Garner et al., 2004). In our facilities, BB frequency was even lower (less than 0.5% out of 500 animals checked). However, all NOS2KO animals exhibited some degree of BB. This phenotype was described in the manufacturer’s webpage (https://www.jax.org/strain/002596 accessed on Oct 11th 2016), and all attempts to get rid of BB in the colony resulted in the loss of the genotype.

One of the few pharmacological options to treat TTM is clomipramine, a tricyclic antidepressant drug, with preferential inhibition properties on serotonin reuptake system (Swedo et al., 1993, 1989). In line with currently available data, our results indicate that repeated clomipramine administration was able to prevent the evolution of BB in NOS2KO. Both males and females were equally responsive to the drug treatment.

TTM is comorbid to several other psychiatric disorders, such as anxiety and obsessive-compulsive disorder (Grant et al., 2016). To investigate this association in NOS2KO, the animals were submitted to the actimeter test, where analyzes both the total activity and the presence of spontaneous repetitive behavior. Our results indicate a decreased activity of NOS2KO associated to an increase in repetitive movements compared to WT animals. Interestingly, clomipramine was effective only in reducing the last parameter, with no effect on total activity. Despite reduced activity, this strain exhibits an antidepressant-like phenotype (Montezuma et al., 2012), and a deficit in extinction of conditioned fear (Lisboa et al., 2015), but no changes in repetitive behaviors was described. However, compulsive-like behaviors have been observed in many transgenic animals. For example, Francis Lee group (Shmelkov et al., 2010) described increased grooming behavior in SLITRK5KO mice. Rodents lacking the adaptor protein SAPAP3 also presented increased grooming, (Welch et al., 2007). Notwithstanding, this protein is responsible for anchoring glutamatergic receptors in the cell membrane and multiple rare missense variants in its gene sequence were found in OCD and TTM suffering patients (Züchner et al., 2009).

The neurobiology of TTM relies on few studies using brain image techniques suggesting a decrease in frontal cortical areas (mainly inferior frontal cortex) associated with an increase in striatal activity (Chamberlain et al., 2009; Fineberg et al., 2010). In NOS2KO mice we observed decreased levels of NMDA subunit NR1 (the binding site for glycine) in prefrontal cortex, associated to an increase of this subunit in striatum. Increased number of neuronal branches was also found in the striatum of NOS2KO mice, in line with a putative increment in this structure activity. It is plausible to consider that both BB and increased repetitive movements found in these animals reflect the increased striatal NR1 and number of neurites. As mentioned before, the cortico-striatal-thamalic circuitry (CSTC) is a series of reverberatory loops that integrate external signals and trigger motor programs in response. One of the integrative structures in this circuitry is striatum, including its multiple subdivisions, and disturbances in particular ‘hubs’ in the CSTC are associated with different pathologies, from Huntington’s disease to obsessive-compulsive disorder, including trichotillomania [for review see (Langen et al., 2011a, 2011b)]. As a proof of concept, the treatment with the NMDA antagonist memantine was equally effective as clomipramine, in preventing the evolution of BB in NOS2KO mice. Taken together, our results suggest a potential therapeutic approach on TTM with compounds acting on the glutamatergic system. In fact, drugs acting on decreasing glutamatergic neurotransmission have been successfully tested in TTM patients successfully. For example, n-acetyl-cysteine was found effective in alleviating TTM symptoms in both double-blind placebo-controlled and case studies (Coric et al., 2007; Grant et al., 2009; Rodrigues-Barata et al., 2012). This compound also exhibited promising results in OCD suffering patients [for review see (Oliver et al., 2015)]. Riluzole, another compound with antiglutamatergic properties, was also found effective in OCD-related disorders (Emamzadehfard et al., 2016).

Information linking NOS2 to putative neuronal developmental process that could lead to malfunctions in CSTC is also scarce. One of the few pieces of evidence about this topic was provided by Arnhold and colleagues (2002). In this article the authors described the role of NOS2 in the early stages of neuronal maturation, depicting a functional part for this enzyme in regulating the cell differentiation and migration in embryonic cortex, although not through the canonical pathway of NO acting on soluble guanylate cyclase. Cultured cells from E14 rat embryo, when incubated with selective NOS2 inhibitors, showed lower number of microtubule-associated protein (MAP2)- labeled neurons. Similarly, our data indicates that silencing NOS2 expression, leads to a reduction in the levels of MAP2 in activity-dependent differentiation of PC12 cells (Banno et al., 2008). The role of NO in activity-dependent differentiation of PC12 cells have been described [for example (Nakagawa et al., 2000)], however implicating the NOS1 isoform as involved in this process. This discrepancy can rely on the culture conditions between the studies. For example, Nakagawa and colleagues used a serum-free medium and 45mM of KCl to differentiate the cells, while we rely on 0.5% FBS and 25mM of KCl. Moreover, in the former study the authors cultivated a low density of PC12 cells (approximately 50 cells/cm2; in 96-well plate), while we used a high density from the beginning of the experiment – 125000 cells/cm2 (in 12-well plate).

Taken together, we propose that increased BB observed in NOS2.KO is a resultant of inefficient inhibition of striatal motor programs by PFC. Supporting this idea, striatum of NOS2KO exhibit abnormal levels of NR1 subunit and neuronal branches. The lack of NOS2 in the early stages of cortical development could impact the maturation of cortical neurons, as evidenced by reduction of MAP2 expression following NOS2 silencing *in vitro* or reduced NR1 levels in PFC, which in turn would lead to the phenotype. The BB phenotype observed in these animals is sensitive to the treatment with clomipramine and memantine, a NMDA antagonist, strengthening its role as a putative model of TTM. Further experiments will be necessary to clarify the role of NOS2 in the development of BB phenotype and its functional consequences in the CSTC; as well as validate its sensitivity to other drugs.

## Acknowledgements

Acknowledgements: this study was supported by grants from Fapesp (2011/02746-4 and 2012/17626-2), CNPq (471382/2011-6) and European Research Council (iPlasticity, # 322742). The authors thank to Eleni T. Gomes (USP), Outi L. Nikkilä and Sulo J. Kolehmainen (UH) for their technical assistance.

